# Cell growth dilutes the cell cycle inhibitor Rb to trigger cell division

**DOI:** 10.1101/470013

**Authors:** Evgeny Zatulovskiy, Daniel F. Berenson, Benjamin R. Topacio, Jan M. Skotheim

## Abstract

Cell size is fundamental to function in different cell types across the human body because it sets the scale of organelle structures, biosynthesis, and surface transport^1,2^. Tiny erythrocytes squeeze through capillaries to transport oxygen, while the million-fold larger oocyte divides without growth to form the ~100 cell pre-implantation embryo. Despite the vast size range across cell types, cells of a given type are typically uniform in size likely because cells are able to accurately couple cell growth to division^3–6^. While some genes whose disruption in mammalian cells affects cell size have been identified, the molecular mechanisms through which cell growth drives cell division have remained elusive^7–12^. Here, we show that cell growth acts to dilute the cell cycle inhibitor Rb to drive cell cycle progression from G1 to S phase in human cells. In contrast, other G1/S regulators remained at nearly constant concentration. Rb is a stable protein that is synthesized during S and G2 phases in an amount that is independent of cell size. Equal partitioning to daughter cells of chromatin bound Rb then ensures that all cells at birth inherit a similar amount of Rb protein. *RB* overexpression increased cell size in tissue culture and a mouse cancer model, while *RB* deletion decreased cell size and removed the inverse correlation between cell size at birth and the duration of G1 phase. Thus, Rb-dilution by cell growth in G1 provides a long-sought cell autonomous molecular mechanism for cell size homeostasis.

---

Phenomenologically, cell size control is achieved through size-dependent changes in cell growth rate or cell cycle progression^5,6,13,14^. Size-dependent changes in cell growth rate can control size when larger cells grow slower than cells nearer the target size, which may be due to an observed optimum cell size for mitochondrial potential^15^. For size-dependent cell cycle progression, larger cells progress through specific phases of the cell cycle more rapidly than smaller cells. Size control is then driven by smaller cells having more time to grow. Evidence for a size-dependent G1/S transition was first provided over 50 years ago^3^ and a series of recent single-cell experiments showed an inverse correlation between cell size at birth and the duration of G1 phase^4–6^. While cell size can be affected by the deletion, overexpression, or chemical inhibition of some cell cycle regulators^7–12,16^, the molecular mechanism through which these genes convert cell size into a biochemical activity driving the G1/S transition is unknown.

One mechanism through which cell size and growth can regulate a cell cycle transition is via the dilution of specific cell cycle inhibitors, while other regulatory proteins remain at a constant concentration^17^ (Fig. 1a). This model was recently shown to apply to budding yeast G1 control, where cell growth diluted the cell cycle inhibitor Whi5 to trigger cell cycle progression^18^. Whi5 inhibits the G1/S transition by binding and inhibiting key downstream transcription factors^19,20^. In mammals, Whi5’s functional orthologs are the pocket protein family comprising the retinoblastoma-associated protein Rb, p107, and p130^21^. The pocket proteins bind and inhibit the activating E2F transcription factors to control the G1/S transition^22,23^ (Fig. 1b). Just as deletion of Whi5 in yeast leads to smaller cells, the deletion of pocket family proteins reduces the cell size of mouse embryonic fibroblasts^9,10^. Thus, the functional similarity between Whi5 and Rb motivated us to test if Rb dilution was a mechanism linking cell growth to cell cycle progression in human cells.

**Figure 1:**
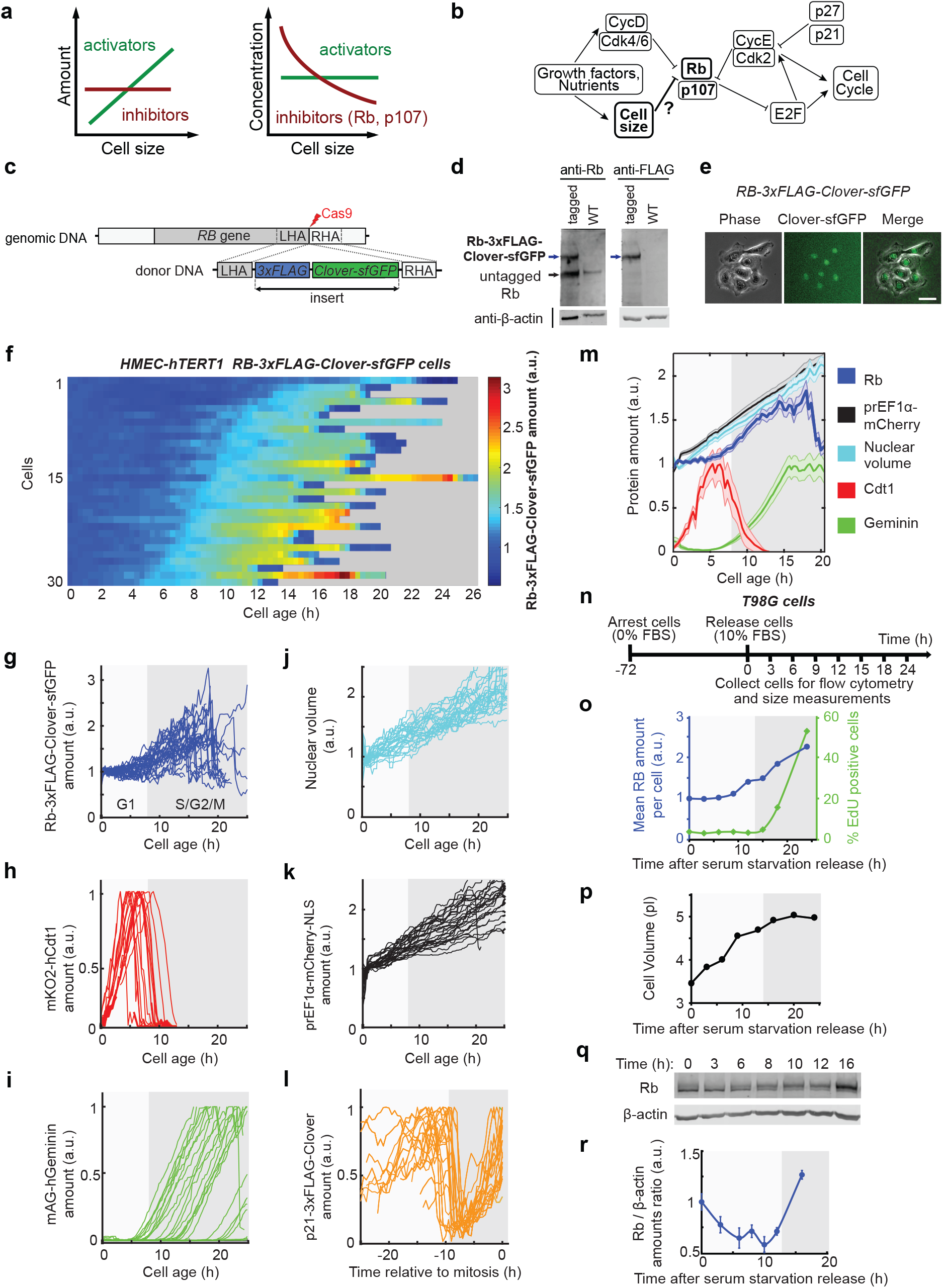
The cell cycle inhibitor Rb is diluted by cell growth in G1. **a**, The inhibitor dilution model: the amounts of key cell cycle activator proteins scale proportionally to cell size, while some cell cycle inhibitors do not to produce a size-dependent signal. **b**, Mammalian G1/S regulatory network. **c**, Endogenous *RB* tagging by CRISPR/Cas9-mediated cleavage and homology directed repair. **d**, Validation of *RB* tagging by immunoblotting with anti-FLAG and anti-Rb antibodies. β-actin is used as a loading control. **e**, Phase, fluorescence, and composite images of HMEC-hTERT1 cells expressing endogenously tagged *RB-3xFLAG-Clover-sfGFP*. **f-k** Single cell traces showing dynamics of Rb-3xFLAG-Clover-sfGFP (f-g), mKO2-hCdt1 (h), mAG-hGeminin (i), nuclear volume (j), mCherry-NLS expressed from an EF1α promoter (k), and p21-eGFP-3xFLAG (l). Rb-3xFLAG-Clover-sfGFP, mCherry-NLS and nuclear volumes for each cell trace are normalized to the their values at cell birth; p21-eGFP-3xFLAG, mKO2-hCdt1 and mAG-hGeminin are normalized to their maximum values. Single cell traces in (f-k) are aligned by cell birth, and in (l) by mitosis. **m**, Average traces for f-l: solid lines show mean values for each time point, and shaded regions denote the standard error of the mean. **n**, Schematic of the time course experiment with synchronized T98G cells (o-r). **o**, Mean Rb amount per cell and percentage of EdU-positive (S-phase) cells are shown for each time point measured with flow cytometry of 50,000 cells. **p**, Corresponding mean cell size dynamics measured with Coulter counter. A biological replicate for n-p is shown in Extended data Figure 3a-b. **q**, Characteristic immunoblots showing the amounts of Rb and β-actin. **r**, Quantification of immunoblots as in (q) measuring the ratio of Rb and β-actin protein amounts (n = 3). For all panels shaded regions denote the median time of the G1/S transition as measured in (h-i).

To test if cell growth diluted Rb in G1, we used CRISPR/Cas9 to C-terminally tag one copy of *RB* at its endogenous locus with two green fluorescent proteins and a triple FLAG epitope tag (Fig. 1c-d). We tagged *RB* in HMEC-hTERT1 cells, a non-transformed human epithelial cell line previously used to identify mutations affecting cell proliferation^24^. We imaged and tracked asynchronously cycling *RB-3xFLAG-Clover-sfGFP* cells over multiple days using an automated wide field microscope (Fig. 1e, Extended data Fig. 1a-b, Supplementary data Movie 1). Consistent with the Rb-dilution model, the amount of Rb was constant for the first 5-10 hours after mitosis, which corresponded to the G1 phase as measured using a FUCCI cell cycle reporter^25^. Rb amount then increased during S/G2/M cell cycle phases (Fig. 1f-i). Meanwhile, the nuclear volume, within which Rb is distributed, increased steadily through the cell cycle (Fig. 1j). To further confirm that HMEC cells grow and accumulate protein during G1, we integrated a construct expressing a nuclear-localized mCherry fluorescent protein from the constitutive *EF1α* promoter and measured its accumulation (Fig. 1 k). Next, we sought to test for the dilution of p21, another important cell cycle inhibitor implicated in G1 control. As for *RB*, we used CRISPR/Cas9 to tag p21 at its endogenous locus (*CDKN1A*) with a green fluorescent protein (Extended data Fig. 2). Unlike Rb, p21 accumulated in G1 and was rapidly degraded at G1/S, consistent with published reports^26–28^ (Fig. 1l; Supplementary data Movie 2). Taken together, these experiments show that Rb is likely diluted relative to other proteins during G1 (Fig. 1m).

To test if Rb is diluted during G1 in different conditions commonly used for cell cycle experiments, we synchronized T98G cells by serum starvation and then followed Rb cell cycle dynamics (Fig. 1n). We used flow cytometry to measure Rb amount and DNA replication in each cell by immunostaining and EdU incorporation respectively. Consistent with the dilution model, Rb amount was constant until accumulation began just before the initiation of DNA replication (Fig. 1o, Extended data Fig. 3a). To confirm cells grow during G1 in this experiment, the corresponding cell size dynamics were measured using a Coulter counter (Fig. 1p, Extended data Fig. 3b). Consistently, the Rb concentration decreased during G1, as measured by immunoblotting (Fig. 1q-r; see Extended data Fig. 3c-e for Rb antibody specificity controls).

To test if Rb was diluted in different cell lines, we turned to immunostaining and flow cytometry because it does not require genome editing and is high-throughput. Cell size was estimated from forward scatter, which correlated with total protein dye measurements (Extended data Fig. 4a-d), and cell cycle phase was determined using DAPI DNA stain. We found that Rb amounts are independent of cell size in G1, while β-actin amounts increased with cell size in four different human cell lines, including primary lung fibroblasts (Fig. 2a-d). To further confirm Rb dilution, we sorted cells into bins with different mean cell sizes and immunoblotted for Rb and β-actin. Consistent with our previous results, we found that Rb concentration decreases during G1 (Extended data Fig. 4e-f).

**Figure 2:**
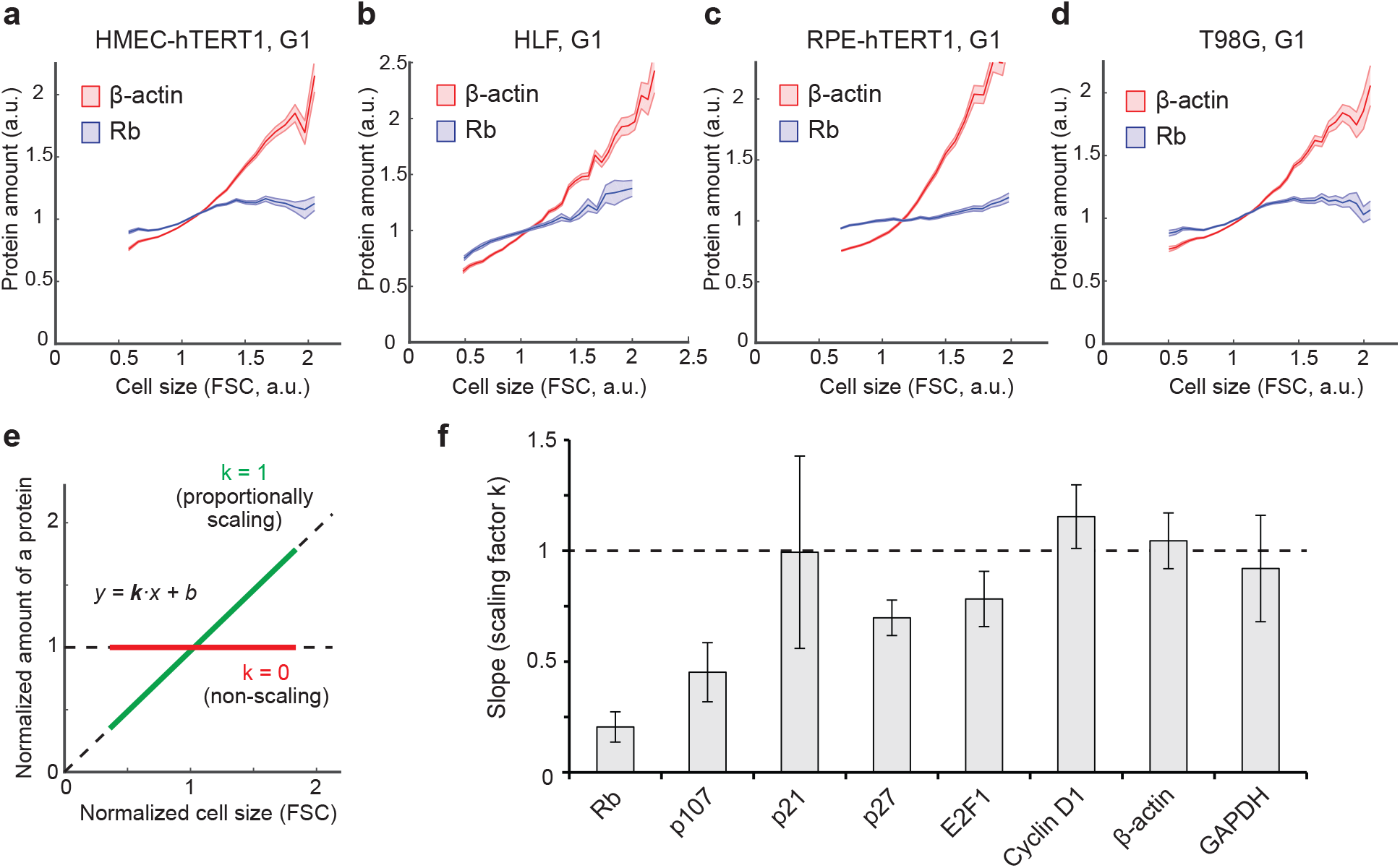
Rb amount in G1 is independent of cell size, while the amount of other G1/S regulators increases with cell size. **a-d**, Amounts of Rb and β-actin proteins in G1 cells of different sizes measured by flow cytometry using HMEC-hTERT1 cells (a), primary human lung fibroblasts, HLF (b), RPE-hTERT 1 (c), T98G (d). Shaded lines show mean values for size bins ± s.e.m. for β-actin (red) and Rb (blue). DNA content was measured with DAPI to identify G1 cells. **e**, A linear fit of cell size and immunofluorescence measurements determine the scaling behavior of the proteins of interest. **f**, Slopes from linear fits for the indicated proteins. Error bars denote s.e.m. shown from biological replicates (n=5). See extended data Fig. 5 for representative flow cytometry data.

Next, we sought to test if other regulator protein amounts increased with cell size during G1 using immunofluorescence and flow cytometry (Extended data Fig. 5a-p). The amount of p107, a close relative of Rb that also inhibits E2F transcription factors, was relatively constant during G1. In contrast, the amounts of the cell cycle activators E2F1 and cyclin D1, and of the cell cycle inhibitors p21 and p27, as well as housekeeping genes such as β-actin and GAPDH, increased with cell size in G1. To characterize the degree of size-dependency of protein amounts, we fit a line to the flow cytometry data after normalizing this data to the mean values (Fig. 2e-f). One mechanism through which an Rb concentration decrease could drive G1/S would be through titration by the amount of activating E2F transcription factors (Extended data Fig. 5q). In this model, larger cells will have higher concentrations of free E2F1 molecules not bound to Rb and will therefore activate transcription to initiate the G1/S transition.

We next sought to determine how Rb dilution in G1 arose from its synthesis and degradation dynamics. To measure *RB* mRNA dynamics, we used fluorescence activated cell sorting to isolate cells from different cell cycle phases and cell size fractions based on their DNA content and forward scatter (Extended data Fig. 6a). From these populations of cells, we then isolated cellular RNA and performed RT-qPCR. *RB* mRNA concentration is low in G1, intermediate in S and high in G2 phase, which reflected our protein measurements (Fig. 1f, 3a). To determine Rb protein stability, we generated an HMEC-hTERT1 cell line carrying a doxycycline-inducible *Clover-3xFLAG-RB* allele (Extended data Fig. 6b-c). We stimulated cells with doxycycline (Dox) for 48h, then washed out Dox and measured Rb amount with immunoblots (Fig. 3b). Rb half-life was ~29 hours, which is longer than the average cell cycle duration 16.6 ± 0.5 hours (± s.e.m.; Fig. 3c, Extended data Fid. 1a), consistent with previous reports that Rb is a stable protein^29,30^. Thus, throughout G1, cells mostly retain the Rb protein inherited from their mother.

**Figure 3:**
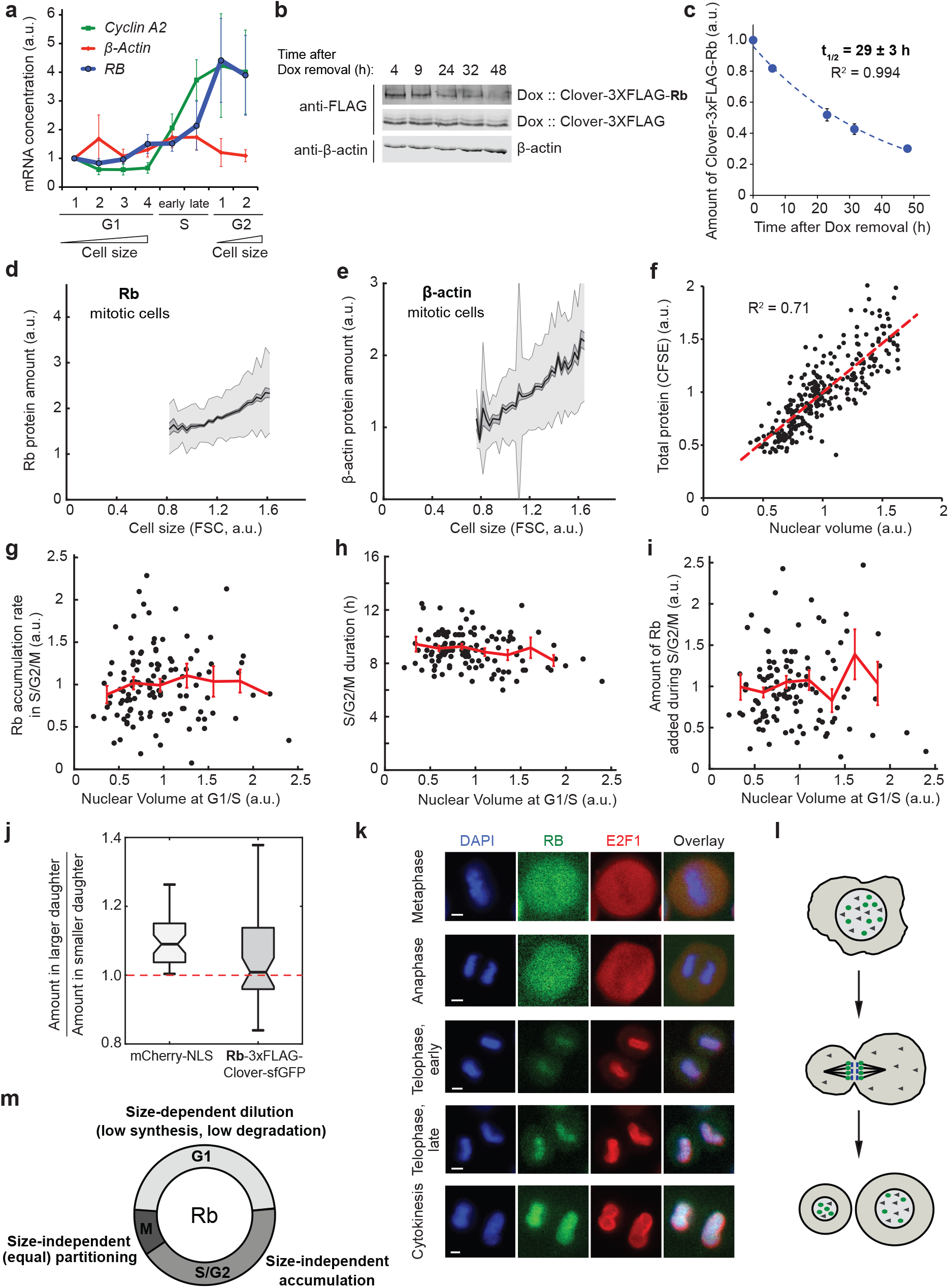
Rb is synthesized size-independently during S/G2 and partitioned equally between daughter cells during mitosis. **a**, Concentrations of *RB, Cyclin A2* and *β-Actin* mRNA measured by RT-qPCR from HMEC-hTERT1 cells sorted by cell cycle phase and cell size using FACS. The values are normalized to *gapdh* mRNA and then scaled by their value in the bin with the smallest G1 cells. Data shown as mean ± s.e.m. (n = 3). **b-c**, Rb protein half-life measurement. Doxycycline-induced *Clover-3xFLAG-RB* expression in HMEC-hTERT1 cells was stopped by doxycycline removal and the amount of Clover-3xFLAG-Rb was measured by immunoblotting (b). Rb half-life was calculated by fitting an exponential decay curve (c). Half-life estimates are ± s.e.m. (n = 3). **d-e**, Amount of Rb (d) and β-actin (e) in mitotic cells as measured by flow cytometry. Mitotic cells were identified by having highly phosphorylated histone H3 (phospho-HH3). Approximately 6,000 mitotic cells were analyzed for each plot. The data were binned by cell size, and mean, s.e.m., and s.d. were plotted for each bin. **f**, Correlation between nuclear volume and CFSE dye, total cellular protein stain, measured by microscopy (n = 284). **g-i**, Rb accumulation rate in S-G2 (g), duration of S-G2 (h) and the total amount of Rb accumulated during S-G2 (i) plotted against nuclear volume at G1/S, n = 122 cells. Red lines show means ± s.e.m. for each nuclear volume bin. **j**, The amount of indicated proteins partitioned to the larger sister cell relative to the amount partitioned to the smaller sister cell, measured with widefield fluorescence microscopy. n = 30 pairs of newborn sister cells. The boxes show lower quartile, median, and upper quartile values. Bars denote the entire range. **k**, Immunofluorescence staining showing the association of Rb with E2F1 and DNA during later stages of mitosis. **l**, A model illustrating size-independent partitioning of a DNA-associated protein during mitosis. **m**, Schematic illustrating Rb dynamics during the cell cycle. In G1, the stable protein Rb is diluted by cell growth, then it is synthesized during S and G2 in a cell size-independent manner, after which it is partitioned equally between daughter cells in mitosis. As a result, newborn cells of different sizes receive similar amounts of Rb.

Cell size control, whereby G1 is shorter for larger-born cells, requires that Rb concentration is lower in larger-born cells. This would be true if larger-born cells inherited from their mothers a similar amount of Rb as smaller-born cells. To test this, we first measured the amount of Rb in mitotic cells identified by their highly phosphorylated histone H3 in flow cytometry (Extended data Fig. 6d). In these mitotic cells, Rb amounts were similar while β-actin amounts increased with cell size (Fig. 3d-e). This size-independence of Rb amounts could be the result of Rb synthesis rate and S/G2/M duration both being independent of cell size. To test this, we imaged live HMEC-hTERT1 *RB-3xFLAG-Clover-sfGFP* cells and estimated cell size as the nuclear volume, which correlates with total protein dye measurements^6^ (see methods; Fig. 3f). Indeed, we found that the synthesis rate of Rb and the duration of S/G2/M are both size-independent (Fig. 3g-i).

Size-independence of Rb amounts at cell birth depends not only on size-independent synthesis in mother cells, but also on the symmetry of partitioning at division. HMEC-hTERT 1 cells partitioned an mCherry-NLS protein reporter so that there was typically a 5-10% difference in amounts between the two daughter cells. However, Clover-3xFLAG-Rb was more accurately partitioned and its median difference was less than 1% (Fig. 3j). This may be due to Rb associating with chromosomes during mitosis, which provides an elegant mechanism for equal partitioning (Fig. 3k-l, Extended data Fig. 6e-f, Supplementary data Movies 3-4). Presumably, this happens because Rb gets dephosphorylated during mitosis^31^, which allows it to associate with DNA-bound E2F through late mitosis and cytokinesis. Thus, through each cell cycle, different-sized cells produce similar amounts of Rb and accurately partition it during division so that newborn cells all inherit a similar amount of Rb (Fig. 3m).

Having shown that Rb is diluted during G1, we sought to determine if the Rb dilution mechanism controls cell size. While some earlier tissue culture studies found a cell cycle and cell size dependency on exogenous Rb expression, others did not^16,32–34^. We therefore sought to first determine if there was an effect of Rb on cell size *in vivo*. To test this, we integrated a vector containing a doxycycline-inducible *Clover-3xFLAG-RB* allele into a Kras^+/G12D^; Trp53^−/−^ mouse pancreatic ductal adenocarcinoma cell line, and then allografted these cell lines into NSG mice by subcutaneous implantation^35^. Tumors were allowed to engraft and grow for five days before we induced expression of the exogenous Rb allele (Fig. 4a). After 14 days of *RB* induction the mice were sacrificed, and the tumors were extracted and analysed by immunohistochemical staining (Fig. 4b). Some tumor cells expressed *Clover-3xFLAG-RB* and some did not. Importantly, cells expressing Clover-3xFLAG-Rb had larger nuclei (Fig. 4c). Since nuclear volume is frequently highly correlated with cell volume^6,36^, this result implies Rb concentration affects cell size *in vivo*.

**Figure 4:**
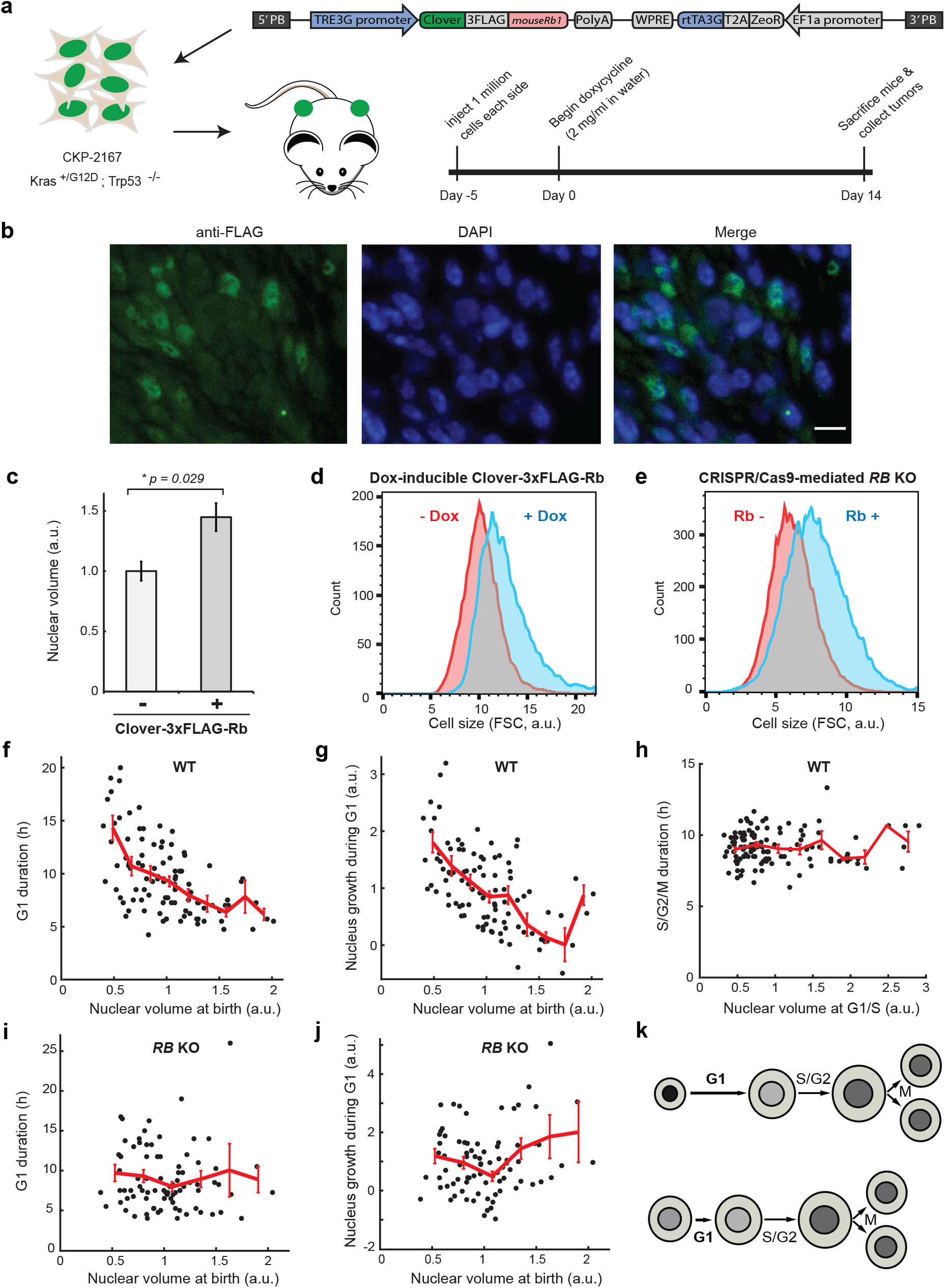
Rb dilution controls cell size. **a**, Experiment schematic showing implantation into mice of tumor cells containing an integrated inducible *Clover-3xFLAG-RB* allele (see text). **b**, Immunohistochemical staining of mouse tumor section with anti-FLAG antibodies 14 days after *Clover-3xFLAG-RB* induction. Anti-FLAG and DAPI nuclear staining are shown in green and blue respectively. **c**, Nuclear volumes of mouse tumor cells expressing or not expressing *Clover-3xFLAG-RB* measured by widefield microscopy. Means ± s.e.m. shown, n = 3 tumors. **d-e**, Flow cytometric size measurements of cultured HMEC-hTERT1 cells carrying a doxycycline-inducible *Clover-3xFLAG-Rb* allele with and without 48 hours of doxycycline induction (d), and of cells with and without *RB* knockout (Rb- and Rb+) (e). **f-g**, G1 duration (f) and the amount of cell growth in G1 (g) as a function of nuclear volume at birth. G1/S timing was defined using the mKO2-hCdt1 FUCCI reporter, n = 110 cells. **h**, S/G2/M duration as a function of nuclear volume at G1/S. **i-j**, *RB* knockout eliminates the inverse correlation between the cell size at birth and G1 phase duration (i) and amount of cell growth during G1 (j), n = 82 cells. 100 nM of palbociclib was added to cell culture media in experiments d-j to partially inhibit cyclin D/Cdk4,6 (see Extended data Figure 8b and corresponding text). **k**, Schematic showing G1 cell size control through the inhibitor dilution mechanism. Shades of grey in the nucleus reflect the concentration of the G1/S inhibitor Rb.

Since Rb affects cell size *in vivo*, we sought to examine a tissue culture model where Rb similarly affects cell size. Such a tissue culture model enables us to measure single cell dynamics to test two key predictions of the Rb dilution model: that size control takes place during the G1 phase of the cell cycle and depends on Rb. We first sought to identify tissue culture conditions that reflect our *in vivo* results, where Rb expression affects cell size. To identify such conditions, we created an *RB* knockout HMEC-hTERT1 cell line and an HMEC-hTERT1 cell line where Rb can be overexpressed with a Dox-inducible promoter. We measured the cell size in both these lines using flow cytometry and compared it to the parental cell line expressing wild-type levels of Rb (Extended data Fig. 7). Indeed, *RB* knockout reduced cell size and Rb overexpression increased cell size. However, the effect was modest (<10%). This relatively small effect may be due to the fact that the vast majority of G1 cells had inactive hyper-phosphorylated Rb, likely due to high cyclin D/Cdk4,6 activity (Extended data Fig. 8a-b), consistent with recently published results^37^. To test if Rb phosphorylation suppressed the effect on cell size that we observed *in vivo*, we generated an HMEC-hTERT1 cell line with a doxycycline-inducible non-phosphorylatable version of Rb where all Cdk sites have been substituted with alanines. Expressing the non-phosphorylatable Rb dramatically increased cell size and the fraction of cells in G1 (Extended Data Fig. 8c-e). We therefore added a low dose (100nM) of the Cdk4,6 inhibitor palbociclib to our media to partially restore Rb activity in wild-type cells (Extended data Fig. 8b). In these conditions, cells continued to cycle, were slightly larger, and were sensitive to changes in Rb concentration (Fig. 4d-e). Based on these results, we conclude that tissue culture cells treated with a low dose of palbociclib provide a good model to investigate Rb-dependent size control.

Size control through size-dependent cell cycle progression manifests as an inverse correlation between the size of a cell when it enters a particular cell cycle phase, and the duration of that cell cycle phase. The increased phase duration then allows smaller cells time to grow more than the initially larger cells. To measure cell size in live asynchronously cycling cells, we used nuclear volume as a proxy because it correlates well with measurements of total protein content (Fig. 3f). Indeed, we observed an inverse correlation between nuclear volume at birth and the duration of G1 as well as the amount of cell growth in G1 (Fig. 4f-g). In contrast, the duration of S/G2/M was not correlated with cell size (Fig. 4h). Moreover, these inverse correlations were eliminated by *RB* knockout (Fig. 4i-j) and much reduced in conditions where Rb concentration had only a small effect on cell size and cell cycle progression (Extended data Fig. 9a-d). This shows that size control takes place during the G1 phase of the cell cycle and depends on Rb (Fig. 4k).

Taken together, our data show that the transcriptional inhibitor Rb and the related protein p107 are diluted by cell growth during the G1 phase of the cell cycle, while other G1/S regulators remain at nearly constant concentration. This provides a long-sought molecular mechanism through which cell growth can drive cell cycle progression in mammalian cells. Importantly, the Rb dilution mechanism does not require progressive phorphorylation by cyclin D/Cdk4,6 in G1 and is therefore consistent with a recent report showing constant hypo-phosphorylation of Rb through most of G1^38^. Finally, we note that Rb-dilution can operate in parallel to other mechanisms, including the recently identified roles for p38 and cell growth rate^5,6,12^.

Our work demonstrates the importance of measuring protein concentration dynamics. Previous genetic studies showed that Rb and p107 deletion, as well as cyclin D or E overexpression, reduced cell size, and genome-wide screens have identified additional regulators involved in size homeostasis including Largen and p38 MAPK^7–12,39^. However, while deletion and overexpression change the concentration of these key regulatory molecules to affect cell size, these experiments do not imply that the concentration of these molecules change in wild type cells as they grow through G1. Our studies of protein concentration dynamics therefore complement the previous genetic studies by identifying the key regulators whose concentrations are in fact changed by cell growth in G1 to drive cell cycle progression, such as Rb and p107.

Conceptually, our work shows how cell size signals can originate in any part of a pathway or network. For a protein to be a ‘size sensor’ all that is required is that the protein’s concentration changes with cell size, and that this concentration change influences cell cycle progression. Here, we identified the size sensors Rb and p107, which are two cell cycle inhibitors in the middle of the mammalian G1/S regulatory pathway. Following similar logic, we previously identified the cell cycle inhibitor Whi5 in budding yeast as a cell size sensor. While Whi5 is functionally similar to Rb, in that both proteins inhibit transcription factors at the core of the G1/S transition, they share no sequence similarity and have a different evolutionary origin^21^. That both these transcriptional inhibitors serve as size sensors by being diluted by cell growth demonstrates a deep conservation of the systems level logic at the core of cell cycle control.

## Materials and Methods

### Cell culture conditions and cell lines

All cells were cultured at 37°C with 5% CO_2_. Non-transformed hTERT1-immortalized human mammary epithelium cells (HMEC-hTERT1) were obtained from Stephen Elledge’s laboratory at Harvard Medical School ^24^ and cultured in MEGM™ Mammary Epithelial Cell Growth Medium (Lonza CC-3150). In microscopy experiments we used the same media but without phenol red to reduce background fluorescence for imaging (Lonza CC-3153 phenol-red free basal media supplemented with growth factors and other components from the Lonza CC-4136 kit). T98G and SaOS-2 cells were purchased from ATCC, recently isolated fetal human lung fibroblasts (HLF) were purchased from Cell Applications, NIH 3T3 cells were obtained from the Rohatgi laboratory at Stanford, RPE-hTERT1 cells were obtained from the Stearns laboratory at Stanford, and CKP-2167 mouse pancreatic ductal adenocarcinoma cells were obtained from the Sage laboratory at Stanford. All these cell lines were grown in Dulbecco’s modification of Eagle’s medium (DMEM) with L-glutamine, 4.5 g/l glucose and sodium pyruvate (Corning), supplemented with 10% FBS (Corning) and 1% penicillin/streptomycin. In cell synchronization experiments T98G cells were kept in DMEM without FBS for 72-96 hours, and then released into the cell cycle by replacing the media with DMEM containing 10% FBS. EdU labeling was performed at each time point by incubating the cells with 10mM EdU for 30 min at 37°C before fixation, staining, and flow cytometry. EdU was detected using a Click-IT EdU detection kit (Thermo Fisher Scientific).

### Fluorescent reporter cell lines

Fluorescent reporters (mKO2-hCdt1, mAG-hGeminin, mCherry-NLS) were cloned into the CSII-EF-MCS lentiviral vector backbone under a constitutive EF1α promoter. The CSII vector, the lentiviral packaging vector dr8.74 and the envelope vector VSVg were introduced into HEK 293T cells by transfection with TurboFect (Life Technologies). Lentivirus-containing medium from transfected HEK 293T cells was used to infect HMEC-hTERT 1 cells. At least 2-3 days after infection, positive cells were sorted by FACS and clonally expanded.

### Rb overexpression and knockout cell lines

For inducible Rb expression, we cloned an Rb expression cassette, driven by the TRE3G doxycycline-inducible promoter (Clontech 631168), into a PiggyBac integration plasmid containing 5’ and 3’ PiggyBac homology arms^35^. The expression cassette contains the human *RB1* or mouse *Rb1* genes fused to fluorescent *Clover* and *3xFLAG* affinity tag sequences, a zeocin resistance gene, and a Tet-On 3G transactivator gene driven by the EF1α promoter. The cell lines stably expressing doxycycline-inducible Rb variants were generated by transfecting cells plated into individual wells of a 6 well plate with 2.2 μg of doxycycline-inducible PiggyBac integration plasmid and 1.1 μg of PiggyBac Transposase plasmid^40^ using the FuGene HD reagent (Promega E2311). Zeocin (400 μg/ml) selection began two days after transfection. Zeocin resistant cells were maintained as polyclonal cell lines.

*RB1* knockout HMEC-hTERT1 cells were generated using the Alt-R® CRISPR-Cas9 system from IDT following the manufacturer’s protocol. Briefly, the guide crRNA:tracrRNA duplexes were formed by mixing a universal tracrRNA with a predesigned specific crRNA in IDT duplex buffer to a final concentration of 3 μM each. For annealing, the mixture was incubated 5 minutes at 95°C and then allowed to cool to room temperature. Then the crRNA:tracrRNA duplexes were mixed with purified Cas9-NLS protein at 1:1 molar ratio in Opti-MEM™ low-serum media (Gibco) for RNP particle formation. The RNP particles were transfected into the cells using the Lipofectamine™ CRISPRMAX™ reagent (Thermo Fisher Scientific). More specifically, for each well in a 6-well plate we added 15 pmol Cas9-NLS, 15 pmol annealed crRNA:tracrRNA duplexes, and 5 μl Cas9 Plus Reagent to 100 μl Opti-MEM™. This mixture was then incubated for 5 minutes at room temperature. We then added 7.5 μl Lipofectamine™ CRISPRMAX™ dissolved in 100 μl Opti-MEM™ and incubated the resulting mixture for 10 minutes at room temperature. The resulting mixture was then added dropwise to each well. We used two different crRNAs that target two sites near the 5’-part of the *RB* coding sequence: GTTCGAGGTGAACCATTAAT (IDT design ID: Hs.Cas9.RB1.1.AA) and AAGTGAACGACATCTCATCT (IDT design ID: Hs.Cas9.RB1.1.AB). In both cases, we obtained about 40-50% knockout efficiency as measured using immunofluorescence staining of Rb and flow cytometry. The resulting cell populations containing both Rb-positive and Rb-negative cells were used for further experiments, where we distinguished these two populations by staining with anti-Rb antibodies.

### Endogenous gene tagging with CRISPR/Cas9

The same CRISPR/Cas9 RNP protocol as for the *RB* knockouts was used, except we used the crRNA targeting the 3’-end of the protein coding sequence to tag proteins at their C-termini (RbC guide: TAGCATGGATACCTCAAACA; p21C: GGAAGCCCTAATCCGCCCAC). We added 4 μg of linearized donor DNA per well to the transfection mixture to introduce the tag by homology directed repair. The donor DNA included left and right homology arms (LHA, RHA) corresponding to the genomic sequences directly upstream and downstream of the Cas9 cut site (for Rb the LHA was 666 b.p long and RHA was 654 b.p. long; for p21 both LHA and RHA were 690 b.p. long). The donor DNA had a synonymous mutation destroying the PAM site, and, between the homology arms, a sequence coding for the chosen tag (2xGGGSG linker-eGFP-3xFLAG for p21, and 3xFLAG-3xGGGSG-Clover-3xGGGSG-sfGFP for Rb). This cassette was followed by a thymidine kinase gene for negative selection against the cells with off-target incorporation of the donor DNA. The cells were cultured in the presence of ganciclovir for negative selection, then GFP-positive cells were sorted clonally by FACS. The clones were expanded and screened using PCR, Sanger sequencing, and immunoblotting to confirm correct tagging.

### Cell imaging

In preparation for imaging, cells were seeded on 35-mm glass-bottom dishes (MatTek) at low density and incubated overnight at 37°C and 5% CO_2_. Then, the cells were moved to a Zeiss Axio Observer Z1 microscope equipped with an incubation chamber and imaged for 48 to 72 hours. Brightfield and fluorescence images were collected from three dishes at multiple positions every 10-20 minutes using an automated stage controlled by the Micro-Manager software^41^. We used a Zyla 5.5 sCMOS camera, which has a large field of view allowing us to track motile cells for many hours, and an A-plan 10x/0.25NA Ph1 objective. The width of the focal plane of this objective is large enough to capture fluorescent protein emissions from the entire nucleus without z-stacks.

### Flow cytometry and cell sorting

For flow cytometry analysis, cells were grown on dishes to ~50% confluence and harvested by trypsinization. The cells were then fixed with 3% formaldehyde for 10 minutes at 37°C and permeabilized with 90% methanol for 30 minutes on ice. Fixed and permeabilized cells were washed once with PBS, blocked with 3% BSA in PBS for 30 minutes at 37°C, and then stained with primary antibodies for 2 hours at 37°C. The cells were then washed twice with a wash buffer (1% BSA in PBS + 0.05% Tween^®^ 20), stained with the fluorophore-conjugated secondary antibodies Alexa Fluor 488 goat anti-mouse (Life Technologies A11029), and/or Alexa Fluor 594 goat anti-rabbit (Life Technologies A11037), and/or Alexa Fluor 405 goat antirat (Abcam ab175673) at 1:1000 dilution for 1 hour at 37°C, and then washed twice again. After this treatment, the cells were resuspended in PBS containing 3 μM DAPI for DNA staining, incubated for 10 minutes at room temperature, and then analysed on a BD LSRII.UV flow cytometer. For plotting, all protein amount and cell size values were normalized to the means for each sample. We performed at least three biological replicates for each experiment.

FACS was used to sort live cells by their size and cell cycle phase. To do this, the cells were harvested from dishes by trypsinization, stained with 20 μM Hoechst 33342 DNA dye in PBS for 15 minutes at 37°C and then sorted on a BD FACSAria flow cytometer. DNA content and the mKO2-hCdt1 fluorescent reporter were used to determine cell cycle phase, and the forward scatter area parameter was used as a readout for cell size. During sorting, cell samples were kept at 4°C, and the sorted cells were collected for further RNA isolation and RT-qPCR analysis or for immunoblotting into corresponding lysis buffers on ice.

### Primary antibodies

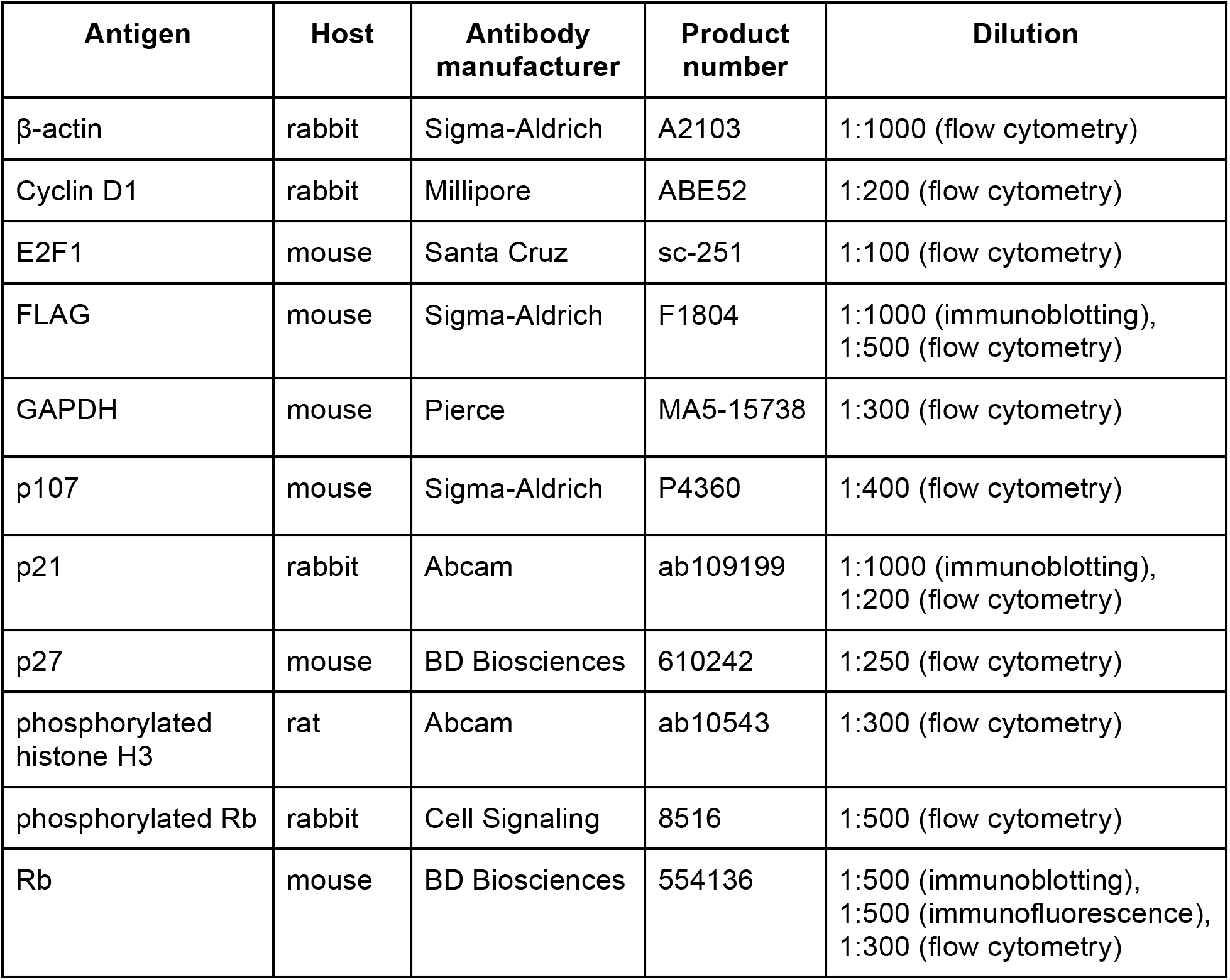

| **Antigen** | **Host** | **Antibody manufacturer** | **Product number** | **Dilution** |
| --- | --- | --- | --- | --- |
| β-actin | rabbit | Sigma-Aldrich | A2103 | 1:1000 (flow cytometry) |
| Cyclin D1 | rabbit | Millipore | ABE52 | 1:200 (flow cytometry) |
| E2F1 | mouse | Santa Cruz | sc-251 | 1:100 (flow cytometry) |
| FLAG | mouse | Sigma-Aldrich | F1804 | 1:1000 (immunoblotting),<br>1:500 (flow cytometry) |
| GAPDH | mouse | Pierce | MA5-15738 | 1:300 (flow cytometry) |
| p107 | mouse | Sigma-Aldrich | P4360 | 1:400 (flow cytometry) |
| p21 | rabbit | Abcam | ab109199 | 1:1000 (immunoblotting),<br>1:200 (flow cytometry) |
| p27 | mouse | BD Biosciences | 610242 | 1:250 (flow cytometry) |
| phosphorylated histone H3 | rat | Abcam | ab10543 | 1:300 (flow cytometry) |
| phosphorylated Rb | rabbit | Cell Signaling | 8516 | 1:500 (flow cytometry) |
| Rb | mouse | BD Biosciences | 554136 | 1:500 (immunoblotting),<br>1:500 (immunofluorescence),<br>1:300 (flow cytometry) |

### Immunoblotting

For immunoblotting, cells were lysed in urea lysis buffer on ice. Proteins from lysates were separated on 10% SDS-PAGE gels and transferred to nitrocellulose membranes. Membranes were then blocked with SuperBlock™ (TBS) Blocking Buffer (Thermo Fisher Scientific) and incubated overnight at 4°C with primary antibodies in 3% BSA solution in PBS. The primary antibodies were detected using the fluorescently labeled secondary antibodies IRDye^®^ 680LT Goat anti-Mouse IgG (LI-COR 926-68020) and IRDye^®^ 800CW Goat anti-Rabbit IgG (LI-COR 926-32211). Membranes were imaged on a LI-COR Odyssey CLx and analyzed with LI-COR Image Studio software.

### RT-qPCR

Total cellular RNA was isolated using an RNeasy kit (QIAGEN). Quantitative RT-PCR (RT-qPCR) was performed using an iTaq Universal SYBR green one-step kit (Bio-Rad) on an iQ5 Bio-Rad instrument. For each sample, the measured mRNA amounts were normalized to *gapdh*, and then for each gene, the resulting mRNA concentration values were scaled by dividing them by their mRNA concentration in the smallest G1 cell sample. RT-qPCR experiments were performed with two technical replicates and at least 3 biological replicates. We used qPCR primers designed by IDT: *gapdh* (Hs.PT.39a.22214836); *β-actin* (Hs.PT.39a.22214847); *RB* (Hs.PT.58.4536695); *Cyclin A2* (Hs.PT.56a.4535284).

### Immunofluorescent staining for microscopy

Cells were seeded on a 35-mm collagen-coated glass-bottom dish (MatTek) one day before immunofluorescent staining. For the staining, cells were fixed with 4% formaldehyde for 10 minutes at room temperature, permeabilized with 0.2% Triton™ X-100 (Sigma-Aldrich) for 15 minutes at 4°C, and then blocked with 3% BSA in PBS. Then the cells were incubated with primary mouse anti-Rb antibodies (BD Pharmingen, 554136) at 1:500 dilution overnight at 4°C, washed twice with PBS, and then incubated with Alexa Fluor 647 conjugated goat anti-mouse secondary antibodies (Invitrogen, A32728) at 1:1000 for 1 hour at room temperature. The cells were washed twice with PBS and incubated with 300 nM DAPI for 5 minutes at room temperature before imaging. *RB* knockout cells were identified by the absence of anti-Rb staining.

### Tumor immunohistochemistry

Tumor sections were obtained from a previous study^35^. Briefly, in that study, Kras^+/G12D^; Trp53^−/−^ mouse pancreatic ductal adenocarcinoma cells CKP-2167 expressing doxycycline-inducible mouse Rb, N-terminally tagged with a triple FLAG epitope and the green fluorescent protein Clover, were allografted by subcutaneous implantation in NSG mice. For each implantation, approximately 1 million cells were suspended in 100 μl of PBS and mixed with 100 μl of Matrigel^®^ Basement Membrane Matrix (Corning 356237). This mixture was injected into the left and right flanks of each mouse. After five days of engraftment and growth, mice were given water supplemented with doxycycline (2 mg/ml) for two weeks. At the end of the experiment, mice were sacrificed, and tumors were harvested, fixed, and paraffin embedded for immunohistochemical analysis.

Immunohistochemical staining was performed on formalin-fixed, paraffin embedded mouse tumor sections. Tumors were fixed in 4% formalin in PBS for 16 hours and stored in 70% ethanol until paraffin embedding. A citrate-based solution (Vector Laboratories) was used for antigen retrieval, and sections were stained with anti-FLAG^®^ M2 monoclonal primary antibodies (Sigma F1804) at 1:500 and subsequently stained with Alexa Fluor 594 conjugated goat antimouse IgG (H+L) secondary antibodies (Invitrogen A-11032) at 1:2000. Nuclei were stained with DAPI. Widefield fluorescence images were collected with a Zeiss Axio Observer Z1 microscope. The cells expressing *Clover-3xFLAG-RB* were identified by positive anti-FLAG staining. The size of cells was estimated by measuring the nuclear area based on DAPI staining and assuming that the nuclear volume is proportional to (Nuclear area)^3/2^. Three tumors from three different mice were analyzed. For every tumor, 160 to 200 Clover-3xFLAG-Rb-positive and Clover-3xFLAG-Rb-negative cells were analyzed from four different fields of view and an average nuclear size was calculated for each individual tumor. Clover-3xFLAG-Rb-positive and Clover-3xFLAG-Rb-negative cells were compared using a two-tailed t-test. Each tumor was then treated as a biological replicate (n = 3).

### Image analysis

For microscopy data quantification, cell nuclei were segmented using the mCherry channel for mCherry-NLS expressing cells or the GFP channel for GFP-tagged Rb and p21 expressing cells. Segmentation was performed by Gaussian filtering and thresholding using manually chosen parameters, followed by opening and closing the segmented regions using the Matlab functions imopen and imclose. Nuclei were manually tracked over time with the aid of a custom Matlab graphical interface. In cases where automated segmentation failed to separate neighboring nuclei, the fused object was split by applying a watershed algorithm while decreasing the morphological structuring element until the desired number of distinct nuclei appeared. Pixel intensities in each channel were summed after applying a location-dependent intensity adjustment to account for our microscope’s optics, where the center of an image is illuminated more brightly than the periphery. Then, each object’s local background was subtracted to account for fluorescence of the cell medium. For the Rb-3xFLAG-Clover-sfGFP and p21-eGFP-3xFLAG cells, we verified the results of this semiautomatic tracking pipeline by manually segmenting and tracking the cells using ImageJ. For each cell, the transition from G1 to S phase was determined by the onset of Geminin accumulation or Cdt1 degradation. To determine the Rb accumulation rate in S/G2, a linear fit was used to approximate the Rb dynamics for each cell, and the the Rb accumulation rate was measured as the slope of this line.

## Acknowledgements

We would like to thank the Elledge, Sage, Stearns and Rohatgi labs for the cell lines used in this study; Dr. Rob de Bruin, Dr. Julien Sage and Dr. Seth Rubin for critical reading of the manuscript; all present and former members of the Skotheim lab for help and valuable discussion, and NIH for funding (GM092925 & GM115479). Cell sorting and flow cytometry analysis for this project were done on instruments in the Stanford Shared FACS Facility: flow cytometry data were collected on the LSRII.UV instrument, and sorting was performed on the Falstaff instrument, both of which were obtained using NIH S10 Shared Instrument Grants (S10RR027431 and S10RR027431, respectively).

## Author contributions

E.Z. and J.M.S. conceived this study, designed and interpreted experiments. D.F.B. generated the prEF1α-mCherry-NLS cell lines and performed and analyzed single-cell tracking experiments with these cell lines, B.R.T. generated doxycycline-inducible cell lines and performed the mouse tumor implantation experiment, E.Z. performed and analyzed all other experiments. E.Z., D.F.B. and J.M.S. wrote the manuscript.

